# Membrane linkage of a *Streptococcus pneumoniae* Wzy capsular polysaccharide occurs through an acylglycerol

**DOI:** 10.1101/2020.09.16.299636

**Authors:** Thomas R. Larson, Janet Yother

## Abstract

Capsular polysaccharides (capsules) protect bacteria from environmental insults and can contribute to virulence in pathogenic bacteria. Their appropriate display on the bacterial surface is critical to their functions. In Gram-positive bacteria, most capsules are synthesized by the Wzy polymerase-dependent pathway, which is also utilized in the synthesis of many capsules and O-antigens of Gram-negative bacteria. Synthesis of capsule repeat units initiates on undecaprenyl-phosphate on the inner face of the cytoplasmic membrane, with polymerization occurring on the outer face of the membrane. In Gram-positive bacteria, the capsule can be transferred to peptidoglycan, as in *Streptococcus pneumoniae* where a direct glycosidic bond to the peptidoglycan *N*-acetylglucosamine occurs. In *S. pneumoniae*, capsule can also be detected on the membrane, and this has generally been assumed to reflect polysaccharide that is linked to undecaprenyl-phosphate and in the process of synthesis. We provide evidence here, however, that final membrane linkage occurs through an acylglycerol, and essentially all of the polysaccharide is transferred from the initial undecaprenyl-phosphate acceptor to an alternate acceptor. This step allows for recycling of undecaprenyl-phosphate and represents an additional terminal step in capsule synthesis. In this regard, capsule synthesis resembles that of the wall- and lipoteichoic acids of *S. pneumoniae*, wherein a common repeat unit and polymer structure are synthesized by the Wzy pathway with divergence at the terminal step that results in linkages to peptidoglycan and a membrane acylglycerol anchor.

**IMPORTANCE:** Linkage of capsular polysaccharides to the bacterial cell surface is a critical step in assuring the ability of these polymers to fulfill their functions, such as the resistance to complement-mediated phagocytosis that can be essential for pathogenic organisms to survive in host environments. Knowledge of the mechanisms by which these linkages occur is incomplete. In this study, we provide evidence for linkage of an *S. pneumoniae* Wzy capsule to an acylglycerol, the most abundant class of lipids in the membrane. This linkage provides a terminal acceptor for capsule that occurs in addition to that of peptidoglycan. Transfer to these terminal receptors is an essential step in CPS synthesis, as failure to do so can be lethal for the cell.

## INTRODUCTION

In pathogenic bacteria, surface polysaccharides provide protection against innate host defenses, particularly those related to complement. In *Streptococcus pneumoniae*, elaboration of a capsular polysaccharide (capsule, CPS) is essential to virulence and colonization. Here, CPS functions through blocking access of phagocytic receptors to complement (C3b) bound to the bacterial cell wall, as well as reducing the deposition of complement, impeding recognition of underlying antigens to antibodies, and limiting mucus-mediated clearance at mucosal sites (1–7). The Wzy pathway for polysaccharide synthesis has been recognized in Gram-positive bacteria, where it is the predominant means of producing capsular polysaccharides and teichoic acids, and in Gram-negative bacteria, where it is a major mechanism for producing capsules and the O-antigens of lipopolysaccharides (8–10). In *S. pneumoniae*, 98 of the described 100 serotypes are synthesized via the Wzy pathway, with the remaining two serotypes being synthesized by the synthase pathway (10–12). The fundamental Wzy pathway is the same in both Gram-positive and Gram-negative bacteria, with variations occurring in sugar contents, linkages, and final destinations on the cell (i.e., outer membrane in Gram-negative bacteria; peptidoglycan and, as shown in the present study, cytoplasmic membrane in Gram-positive bacteria,). For *S. pneumoniae* CPS serotype 2 (Fig. 1), synthesis begins on the inner face of the cytoplasmic membrane with the CpsE-catalyzed transfer of β-glucose-1-phosphate (Glc-1-P) from UDP-glucose (UDP-Glc) to the C_55_ lipid undecaprenyl-phosphate (Und-P) (13). The additions of subsequent sugars are catalyzed by the glycosyltransferases CpsT (rhamnose, Rha), CpsF (two Rha), CpsG (Glc), and CpsI (glucuronic acid, GlcA) (14, 15). The Wzx flippase translocates the lipid-linked oligosaccharide repeat unit to the outer face of the cytoplasmic membrane. There, the Wzy polymerase mediates synthesis of high molecular weight polysaccharide by linkage of the reducing end of a polymer chain to a nascent repeat unit, with subsequent recycling of the Und-P lipid acceptor. The CPS can then, in varying proportions, remain attached to the membrane, be transferred to the peptidoglycan, or be released into the surrounding milieu (16, 17). In Gram-negative bacteria, the capsule or O-antigen is further transported across and linked to the outer membrane (18, 19).

**Fig 1.**
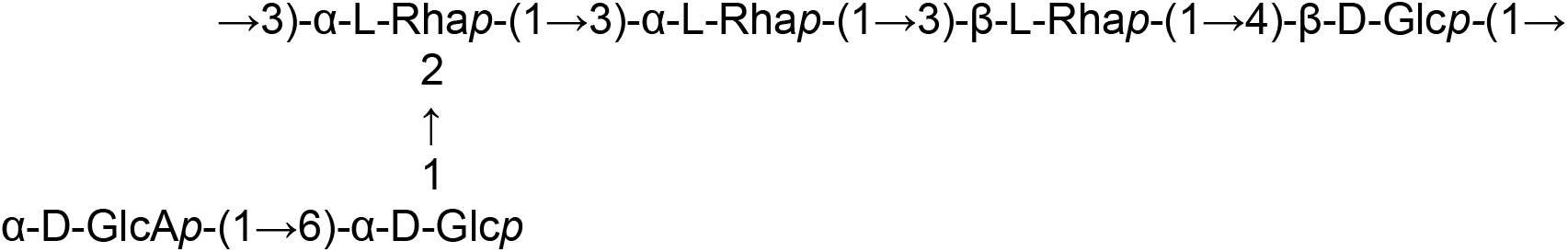
Biological repeat unit of *S. pneumoniae* CPS serotype 2. Initiation of repeat unit synthesis and linkage of CPS to peptidoglycan occur via the β-D-Glc*p-*(1→ of the linear backbone (13, 20). Glc, glucose; GlcA, glucuronic acid; Rha, rhamnose.

In *S. pneumoniae*, the completed CPS polymer exhibits a broad range in size, from only a few repeat units to as many as 2000 repeat units (16, 20, 21). Similar, if not identical, size distributions of polymer are seen on membrane and peptidoglycan fractions for multiple serotypes (16), suggesting that transfer may be random and lacking in a requirement for a termination step that requires a long-chain polymer. Unlike the linkages of other polysaccharides to peptidoglycan, where a phosphodiester bond is involved (22–28), linkage of the serotype 2 CPS to peptidoglycan occurs via a direct 1,6 glycosidic linkage between the reducing end Glc of CPS and the β-D-*N*-acetylglucosamine (GlcNAc) of peptidoglycan (Fig. 1) (20). The serotype 8 and 31 capsules, whose reducing-end sugars are Glc and galactose respectively, are also attached to peptidoglycan through direct glycosidic linkages (20). There have been no published reports suggestive of CPS transfer from Und-P to an alternate membrane acceptor, and it is generally assumed that capsule detected on the membrane is still in the active process of synthesis prior to transfer to peptidoglycan or release from the cell (the latter of which also lacks a description of its mechanism). However, several pieces of data led us to propose that the majority of membrane-bound capsule is not linked to Und-P (17).

In *S. pneumoniae*, high amounts of CPS are detectable on both the membrane and peptidoglycan (16, 17). However, the phenotypes of certain mutants suggested circumstances in which high levels of membrane-associated CPS are lethal for the bacteria. Specifically, mutations that result in the inability to complete CPS synthesis once the addition of the first Rha has occurred (the committed step, catalyzed by CpsT) result in lethality (14, 15, 17). Isolates containing such mutations are obtained only as a result of spontaneous secondary suppressor mutations that result in reduced or no CPS synthesis, which allows for survival (14, 15, 17). In our studies, the primary mutations were generated in *cpsK* (encoding the UDP-glucose dehydrogenase responsible for catalyzing synthesis of UDP-GlcA, the donor for the terminal side chain sugar, Fig. 1), *wzx* (flippase), *wzy* (polymerase), and genes encoding the glycosyltransferases CpsF, CpsG, and CpsI. A requirement for secondary mutations was first suggested for *cpsK* mutants when repair of the primary mutation failed to restore full capsule levels (17). More than 90% of the secondary mutations occurred in *cpsE* (14, 17), which encodes the initiating glycosyltransferase (13), and resulted in a CpsE with greatly reduced or no activity (17). Subsequent analyses demonstrated similar secondary mutations for mutants with each of the other primary mutations mentioned above (14, 15, 17).

For *cpsK* deletion mutants, low amounts of high molecular weight CPS are detected on the membrane fraction but no CPS is detected on the cell wall, indicating an inability to transfer the CPS. Lethality resulting from the primary *cpsK* mutations is expected to be due to the accumulation of Und-P linked intermediates that either destabilize the membrane or sequester Und-P such that it is no longer recycled for use in the essential peptidoglycan pathway. Because there are only low levels of Und-P in the cell relative to other lipids (29, 30), such sequestration would result in lethality, an outcome that has also been suggested for mutations affecting Wzy polysaccharide synthesis in other bacteria (31–34).

The low levels of membrane-linked CPS in *cpsK* mutants may result from the limited amount of Und-P utilized in the CPS pathway due to reduced CpsE activity, and/or the amount of Und-P linked intermediates that can accumulate in the membrane without detriment to the cell. However, the latter interpretation is inconsistent with the large amounts of membrane-linked CPS that accumulate in parental strains without harm (16, 17). We therefore hypothesized the existence of an alternate membrane acceptor, with lethality in the mutants resulting from the failure to transfer repeat units from Und-P to either peptidoglycan or this acceptor. In the present study, we examined the mechanism of CPS-membrane linkage using chemical, enzymatic, and composition analyses via gas chromatography – mass spectrometry (GC–MS). The results demonstrate the membrane-associated CPS is linked to an acylglycerol carrier. Thus, while CPS synthesis initiates on Und-P, subsequent steps result in its transfer and retention on the cell by linkage to either peptidoglycan or an alternate lipid acceptor in the membrane.

## RESULTS

### Linkage of CPS to the membrane does not involve a polyprenyl- or glycerophosphate anchor

In a previous study analyzing linkage of CPS to peptidoglycan, metabolic labeling of growing bacteria did not reveal incorporation of ^32^P into high-molecular-weight, membrane-associated CPS (20), possibly indicating phosphate bonds are not involved in the anchoring. To further test this result, we purified *S. pneumoniae* CPS-containing membranes from the serotype 2 strain D39 and examined the susceptibility of the bond to chemical and enzymatic treatments.

The pyrophosphate bonds of Und-P-P-linked sugars and polysaccharides are highly sensitive to mild acid hydrolysis (35). In *in vitro* experiments, Und-P-P-Glc synthesized using CpsE-containing membranes and exogenous UDP-Glc is completely cleaved by mild acid hydrolysis to liberate Glc (13). However, similar treatment of isolated membranes containing *in vivo* synthesized CPS released only a small amount of polymer (Fig. 2). This released CPS may represent Und-P-P-CPS that exists prior to transfer to peptidoglycan or another acceptor.

**Fig. 2.**
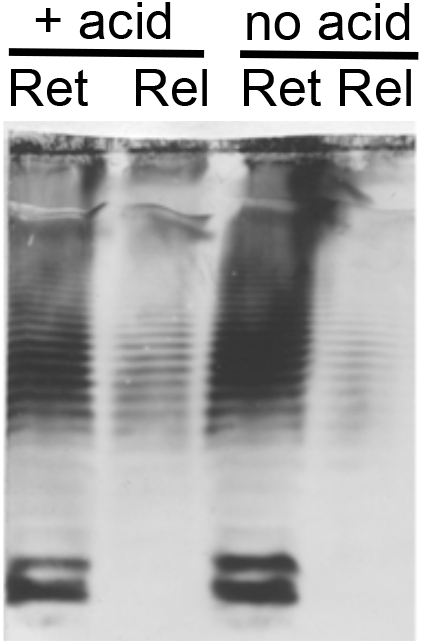
CPS-membrane linkages are largely resistant to mild-acid treatment. Isolated membranes from the serotype 2 D39 strain were incubated at 70°C for 20 min in reactions containing 0.1 M HCl (+ acid) or 0.1 M KCL (no acid). The reactions were then neutralized with KOH or brought to a salt concentration of 0.2 M with KCl, respectively. Following centrifugation (135,000 ×*g* for 30 min at 4°C), the CPS retained on the pelleted membranes (Ret) and released into the supernatant (Rel) was examined by SDS-PAGE and immunoblotting with serotype 2-specific antiserum.

Phospholipase D (PLD) is a phosphodiesterase that cleaves the phosphodiester bonds in glycerophospholipids and in glycerophospholipid-linked polysaccharides. Treatment with PLD did not release CPS from membranes of either the serotype 2 D39 parent strain or an isogenic derivative of D39 that produces only short chain CPS due to deletion of *cpsABCD* (Fig. 3). These genes encode proteins involved in modulation of CPS synthesis, but the mutant strain is identical to the parent in its ability to transfer CPS to the membrane and peptidoglycan (16) (Gupta, Ambrose, Larson, and Yother, manuscript in preparation). Phospholipase A2, phospholipase C, and PLD from a different source also failed to release the serotype 2 CPS (data not shown).

**Fig. 3.**
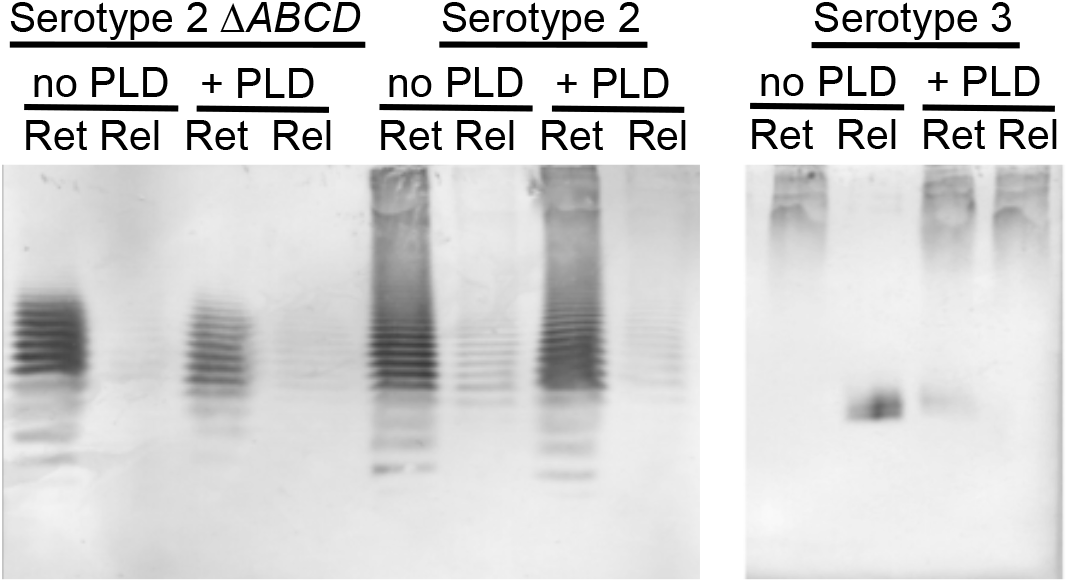
Serotype 2 CPS but not serotype 3 CPS is retained on the membrane following phospholipase D (PLD) treatment. Isolated membranes from serotypes 2 and 3 were incubated with buffer only (50 mM Tris (pH 8) and 10 mM CaCl_2_, no PLD) or with buffer plus PLD at 30°C for 1 hr. Following centrifugation of the reaction mixtures (135,000 × *g* for 30 min at 4°C), the CPS retained on the pelleted membranes (Ret) and released into the supernatant (Rel) was examined by SDS-PAGE and immunoblotting with serotype 2- or serotype 3-specific antiserum. Serotype 2 is the parent D39. Serotype 2 Δ*ABCD* is an isogenic derivative of D39 that produces only short chain CPS. Serotype 3 CPS is produced by a processive mechanism rather than by single repeat unit addition, resulting in only high MW CPS and no ladder formation (16, 36, 38, 39). See text for explanation of the basis for the retained and released serotype 3 CPS. The origin of the low MW species in the released fraction of the no PLD of the serotype 3 CPS has not been determined.

In contrast to the results with the serotype 2 strains, release was observed for the glycerophospholipid-linked serotype 3 CPS (Fig. 3). Synthesis of serotype 3 occurs by the processive synthase-dependent pathway that results in only high MW CPS, as opposed to the single repeat unit addition of the non-processive Wzy pathway (10, 16, 36–39). Serotype 3 CPS is not linked to the peptidoglycan (40). The partial release of CPS following PLD-treatment (Fig. 3) reflects the two distinct CPS populations that occur during synthesis: one that is still linked to the phosphatidylglycerol on which synthesis initiated, and one that has been cleaved from the phosphatidylglycerol but remains cell-associated through engagement with the synthase (10, 39, 41, 42). The former is released by PLD-treatment whereas the latter is not. The observation of PLD-mediated release of serotype 3 CPS from isolated membranes is consistent with previous results demonstrating its release by PLD treatment after *in vitro* synthesis (39, 41, 42) and confirms the same linkage occurs *in vivo*.

### The CPS-membrane linkage is susceptible to mild alkaline treatment

Glycosyl-diacylglycerols are the major lipid component of *S. pneumoniae* membranes (43, 44). These structures are resistant to mild acid but are susceptible to hydrolysis of the acyl chains under mild alkaline conditions, whereas the opposite is true for polyprenol phosphate-linked sugars (35, 45). To determine whether CPS could be released by mild alkaline treatment, purified membranes from the D39 Δ*cpsABCD* strain were used, as the short CPS chains produced by this strain facilitated subsequent studies requiring purification of CPS linked to its membrane anchor (see next section). As observed with the D39 parental strain, the amount of CPS retained on the Δ*cpsABCD* membranes was largely unaffected by treatment with mild acid (Fig 4). In contrast, mild alkaline treatment resulted in a near-complete loss of membrane-associated CPS (Fig 4). Increasingly harsh alkaline conditions showed further reduction in the retained CPS (Fig 4). For both alkaline conditions, only minimal amounts of released products were detected, and the total amount of CPS in membrane and released fractions was less than that for the control and mild-acid treated samples.

**Fig. 4.**
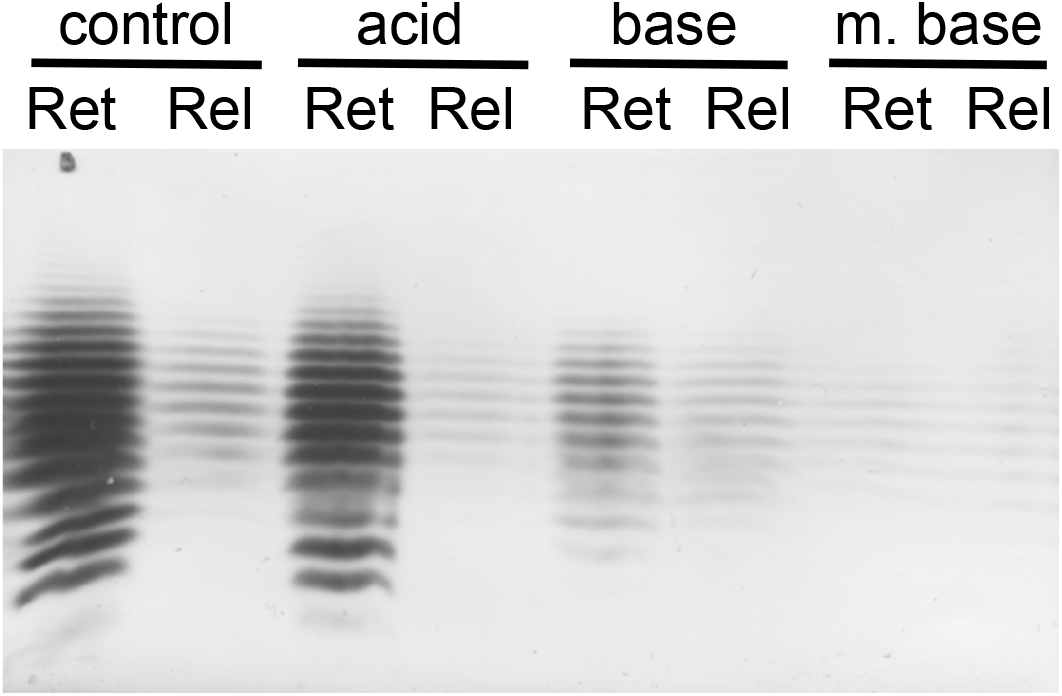
Alkaline treatment removes CPS from the membrane. Isolated membranes from the CPS short chain-producing D39 Δ*cpsABCD* were incubated with 0.1 M KCl (control), mild acid (acid; 0.1 M HCl, 70°C, 20 min), mild alkaline aqueous base (base; 0.1 N NaOH, room temperature, 30 min), or harsher alkaline treatment using methanolic base (m. base; 0.1 N NaOH, 60% methanol, room temperature, 30 min). Following centrifugation of the reaction mixtures (135,000 × *g* for 30 min at 4°C), the CPS retained on the pelleted membranes (Ret) and released into the supernatant (Rel) was examined by SDS-PAGE and immunoblotting with CPS serotype 2-specific antiserum. Lack of CPS in the released fractions of the base conditions is due to degradation by alkaline peeling (see text and Fig. 5).

Alkaline hydrolysis conditions can depolymerize polysaccharides, with sequential degradation and release of single sugars from a free reducing end. This “peeling” reaction can be slowed by the addition of a reducing agent (46–48). To reduce any freed sugars and arrest the alkaline peeling process, the alkaline conditions described above were repeated with the addition of sodium borohydride (NaBH_4_). GC–MS analyses demonstrated loss from the membranes of most of the CPS sugars Rha and Glc under the alkaline plus NaBH_4_ condition in comparison to the control reaction with NaBH_4_ only (Fig. 5A). The CPS sugars from the alkaline plus NaBH_4_ reaction but not the control reaction were detected in the released fraction (Fig. 5B). Similarly, ribitol from the *S. pneumoniae* lipoteichoic acid (LTA) was lost from the membrane under the alkaline plus NaBH_4_ condition (Fig. 5A), with its appearance in the released fraction (Fig.5B). LTA is anchored to the membrane via α-D-glucopyranosyl-(1,3)-diacylglycerol (49), which is one of the two most abundant glycolipids in *S. pneumoniae* membranes (43, 44).

**Fig. 5.**
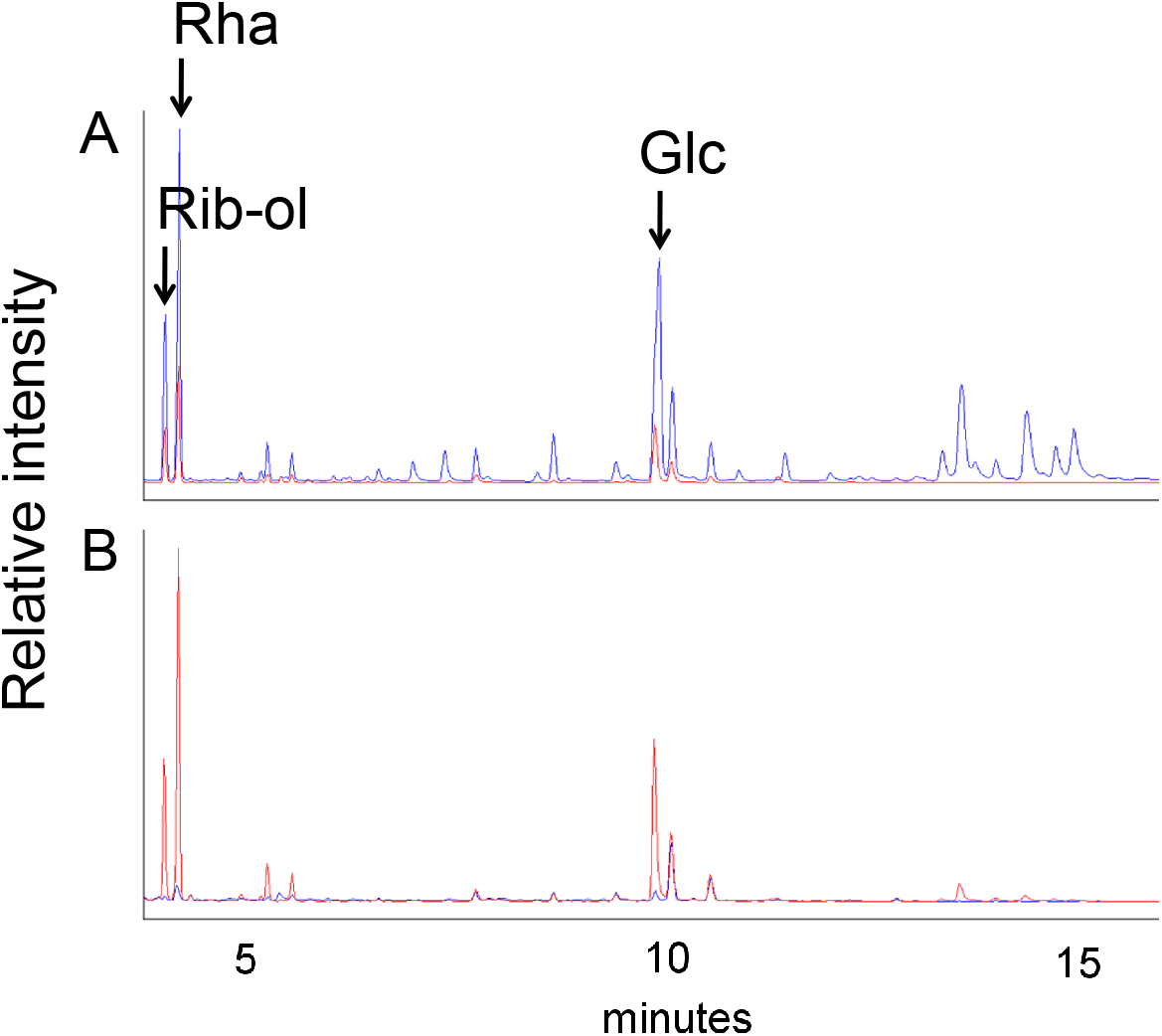
CPS sugars released by alkaline treatment of D39 Δ *cpsABCD* membranes were detected under reducing conditions. Membranes were treated with mild alkali (0.1 N NaOH, room temperature, 30 min) plus the reducing agent NaBH_4_, or with NaBH_4_ alone (control). Following centrifugation of the reaction mixtures (135,000 × *g* for 30 min at 4°C), the products retained on the membrane, and those released into the supernatant, were analyzed by GC–MS. (A) Chromatogram of NaBH_4_-treated control membranes (blue) overlaid with chromatogram of alkali plus NaBH_4_-treated membranes (red) shows significant loss of CPS sugars rhamnose (Rha) and glucose (Glc), as well as the LTA sugar ribitol (rib-ol) from the alkali-treated membranes (red). (B) Overlaid chromatograms of the corresponding supernatants show the CPS and LTA sugars released from the alkali plus NaBH_4_-treated membranes (red), but the near-complete absence of released sugars from the NaBH_4_-treated control membranes (blue). Sugars were identified based on purified standards.

The results obtained by addition of NaBH_4_ to the alkaline reaction (Fig. 5) confirmed that alkaline-induced loss of CPS from the membranes without concomitant detection in the released fraction observed using immunoblotting (Fig. 4) did, in fact, result from release, with subsequent degradation of polymer that resulted in products too small for detection by immunoblotting.

### Isolated lipid-linked CPS contains glycerol

To confirm the acylglycerol linkage suggested by the alkaline hydrolysis results, a Bligh-Dyer extraction (50) was used to further purify lipid-linked CPS from isolated membranes. Here, the CPS-lipid was partitioned into the aqueous-methanol fraction due to the high proportion of CPS relative to lipid. The D39 Δ*cpsABCD* strain was used to facilitate composition analysis, as the shorter CPS chains result in reduced amounts of total CPS sugars in relation to the single glycerol per chain expected for an acylglycerol anchor. The extracted products were purified by anion exchange chromatography to isolate CPS and linked components. The fractions containing CPS were identified by GC–MS analysis for component sugars. Purified CPS contained Rha, Glc, and GlcA in proper ratios expected from CPS repeat units (3:2:1) (Fig 6). The lipid-CPS fraction also contained glycerol, which would be liberated under the methanolysis conditions used to prepare the samples for GC–MS analysis. This result demonstrated that CPS is linked to the membrane through an acylglycerol.

**Fig. 6.**
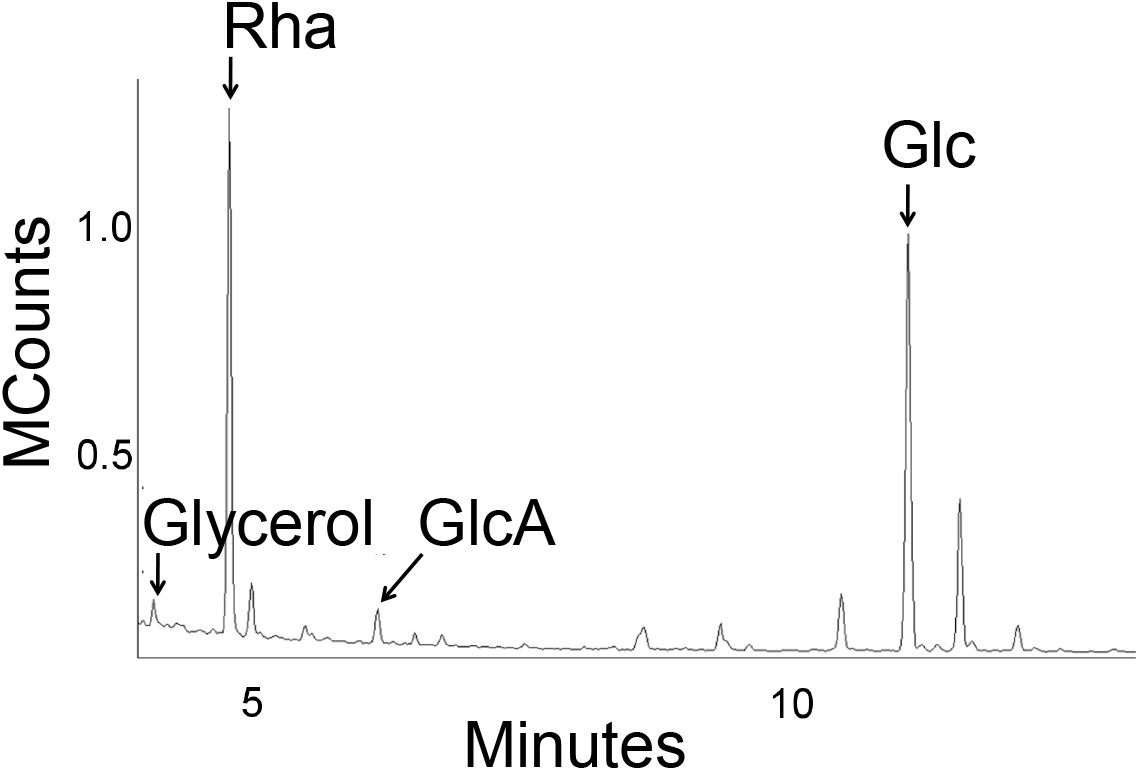
Lipid–linked CPS contains glycerol. The Bligh-Dyer method (50) was used to extract lipid-linked CPS from D39 Δ*cpsABCD*, which was then purified and subjected to GC–MS analysis. In addition to the CPS sugars rhamnose (Rha), glucose (Glc), and glucuronic acid (GlcA), glycerol was identified, consistent with CPS being linked to an acylglycerol. Rha, Glc, and GlcA were present in the ratio expected for CPS repeat units (3:2:1), as determined using purified sugar standards.

## DISCUSSION

Peptidoglycan has been long been known as a site for covalent linkage of capsules synthesized by the Wzy pathway in Gram-positive bacteria (22, 40). Our previous studies demonstrated this linkage in *S. pneumoniae* is a direct glycosidic bond between the reducing end Glc of CPS and the GlcNAc of peptidoglycan (20), unlike the phosphodiester-mediated linkages observed with other polysaccharides (22–28). In addition to the peptidoglycan–localized CPS, we have also reported the presence of a large amount of CPS on the membrane, with our results suggesting this CPS may be linked to an acceptor other than Und-P (16, 17). The results reported in the present study support that outcome and demonstrate that both peptidoglycan and a membrane acylglycerol can serve as terminal acceptors for CPS. Like the CPS-peptidoglycan linkage, the CPS-membrane linkage was not susceptible to mild acid hydrolysis (this study), ^32^P was not incorporated into high molecular polymer (20), and the membrane linkage was additionally resistant to PLD (this study), all of which are properties indicative of a lack of phosphate bonds. The CPS-membrane linkage was susceptible to mild alkaline hydrolysis, with the release of glycerol and the CPS sugars, consistent with cleavage of an acylglycerol membrane anchor.

The finding of two cell-surface terminal receptors for CPS mirrors the mechanisms described for the teichoic acids of *S. pneumoniae*. These polymers are unusual among teichoic acids in that the membrane-linked lipoteichoic acid (LTA) and peptidoglycan-linked wall-teichoic acid (WTA) have the same repeat unit. For both structures, the reducing-end sugar linking them to their respective anchors is AATGal (2-acetamido-4-amino-2,4,6-trideoxygalactose), albeit with different anomeric conformations via a phosphodiester bond for WTA and a glycosidic bond for LTA (23, 49, 51). Like CPS and peptidoglycan, repeat unit synthesis of the teichoic acids initiates on Und-P and follows a Wzy pathway (51, 52). The teichoic acid pathways are common through the point of polymerization on the outer face of the cytoplasmic membrane to achieve a final chain length of 6-8 repeat units (23, 49, 51). Then, the pathways diverge by way of linkage via AATGal-1-P to the peptidoglycan *N-*acetylmuramic acid (MurNAc) residue to yield WTA, or direct linkage via AATGal to α-D-glucopyranosyl-(1,3)-diacylglycerol to yield LTA (23, 49). The latter, along with α-D-galactopyranosyl-(1,2)-α-D-glucopyranosyl-(1,3)-diacylglycerol are the two most abundant glycolipids in *S. pneumoniae* membranes, where they comprise ~30% of the total cellular lipid content (43, 44, 53). This contrasts with other Gram-positive bacteria, where glycosyl-diacylglycerols constitute only 1–2% of the total lipids (54). Glycosyl-diacylglycerols are susceptible to hydrolysis of the acyl chains under mild alkaline conditions. With this treatment, we observed release from the membranes of CPS and LTA. The CPS sugars Glc, Rha, and GlcA, along with glycerol, were detected in the released product but galactose was absent, suggesting that CPS, like LTA, may be linked to the membrane via glucosyl-diacylglycerols. The abundance of these glycolipids in *S. pneumoniae* is consistent with the ability to serve as acceptors for the large amount of CPS detected in the membrane, as opposed to the relatively low concentration of Und-P that would be available (29, 30). The original reports describing the *S. pneumoniae* glycolipids postulated their possible roles in polysaccharide synthesis (55).

The enzymes catalyzing the WTA and LTA transfers to their respective peptidoglycan and membrane receptors are proposed to be LytR and the teichoic acid ligase TacL (51, 56, 57), respectively. The enzyme(s) for CPS linkage have not been established. Deletions of *tacL* do not alter CPS levels, suggesting this enzyme does not perform the dual function of ligating both LTA and CPS to a membrane acceptor (51). It has been suggested that CpsA, the first protein encoded in the *S. pneumoniae cps* locus and a member of the LytR-CpsA-Psr family is involved in linkage to peptidoglycan (58, 59). This family of enzymes was reported to bind polyprenol pyrophosphate lipids and to be phosphotransferases (58), an activity consistent with formation of the phosphate bond in teichoic acid but inconsistent with the direct glycosidic linkage in CPS (20). Additionally, in multiple studies, the attachment of CPS to the peptidoglycan was not altered in Δ*cpsA* mutants or Δ*cpsABCD* mutants (16, 58, 60, 61) (Gupta, Ambrose, Larson, and Yother, manuscript in preparation). Similarly, linkage to the membrane was unaffected in Δ*cpsA* or Δ*cpsABCD* mutants (16) (this study and Gupta, Ambrose, Larson, and Yother, manuscript in preparation).

The results of the present study describe a novel step in CPS synthesis in *S. pneumoniae* and will allow for the complete determination of the membrane anchor and the mechanisms involved in generation of the CPS-membrane linkage. Targeting the terminal steps in CPS synthesis has the potential to attenuate *S. pneumoniae* virulence through both reductions in CPS and lethality to the cell. It remains to be seen whether a similar effect occurs in the CPS of other Gram-positive bacteria, particularly given their low level of glycosyl-diacylglycerols in comparison to *S. pneumoniae*.

## MATERIALS AND METHODS

### Bacterial strains and growth conditions

The *S. pneumoniae* serotype 2 parent D39 (62) and its short-chain CPS derivative *S. pneumoniae* KG920 (D39 Δ*cpsABCD*) ((16) Gupta, Ambrose, Larson, and Yother, manuscript in preparation) were grown statically at 37°C in a chemically defined medium (63). *S. pneumoniae* WU2 (64), a serotype 3 strain, was grown statically at 37°C in Todd-Hewitt Broth (Difco) plus 0.5% Yeast Extract (Difco).

### Isolation of CPS-linked membranes

Membranes were prepared from mid-exponential phase cultures (~3 × 10^8^ CFU/mL) essentially as described previously (13, 16, 36). Briefly, bacteria from 1500 mL cultures were harvested by centrifugation at 20,000 × *g* for 15 min at 4°C, suspended in phosphate buffered saline (PBS; 137 mM NaCl, 2.7 mM KCl, 5.4 mM disodium phosphate, 1.8 mM monopotassium phosphate, pH 7.4), and centrifuged as above. The wash step was repeated. The cell pellet was concentrated 100-fold from the original culture volume in a buffer (20% sucrose, 50 mM MgSO_4_, 50 mM Tris [pH 7.4]) isotonic to the bacterial cytoplasm to maintain protoplasts generated in this step. Forty units per ml of mutanolysin (Sigma) and 2 μl/mL of purified recombinant *S. pneumoniae* autolysin (LytA) (20) were added and the cell suspension was incubated at room temperature overnight. These enzymes release the cell wall by cleaving the peptidoglycan backbone between *N*-acetylmuramic acid and *N*-acetylglucosamine, and removing the peptide side chain from *N*-acetylmuramic acid, respectively. The formation of protoplasts was confirmed microscopically. The protoplasts were harvested by centrifugation at 16,000 × *g* for 20 min at 4°C, washed in the above isotonic buffer, and centrifuged again. The protoplasts were lysed by addition of Milli-Q ultrapure water at 1/100 of the original culture volume, and a short sonication was performed before centrifugation at 135,000 × *g* for 30 min at 4°C. The pelleted membranes were washed with water two additional times and then suspended in Milli-Q ultrapure water at 1/1000 of the original culture volume. After suspension, membranes were kept on ice until use or stored at 4°C for no more than five days. Lack of contamination of membranes with peptidoglycan fragments was confirmed by assay for the peptidoglycan-linked β-galactosidase, as described previously (16).

### Mild acid hydrolysis

Two-hundred fifty μl of membranes isolated as above were treated with acid by the addition of HCl to 0.1 M and water to a final volume of 500 μl. The reactions were incubated at 70°C for 20 min with subsequent neutralization by the addition of dilute KOH. A control fraction consisted of an equal amount of membrane with KCl added to 0.1 M concentration and incubated at 70°C for 20 min. After the incubation, KCl was added to the control reaction to bring the final salt concentration to 0.2 M. The samples were centrifuged at 135,000 × *g* for 30 min at 4°C. The supernatant containing any released CPS was removed and saved, and the membrane pellet was brought to a volume equal to that of the supernatant.

### Phospholipase D treatment

Membranes isolated as above were treated with PLD as previously described (39). Briefly, 500-μl reactions containing two hundred fifty μl of membranes, Tris-HCl (pH 8; final 50 mM concentration), CaCl_2_ (10 mM final concentration) and PLD (25 U/ml, final concentration) were incubated at 30°C for 1 hr. The reactions were then heated at 100°C for 4 min, and the precipitate was removed by centrifugation at 10,000 × *g* for 5 min at 4°C. The resulting supernatants were centrifuged at 135,000 × *g* for 30 min at 4°C. The supernatants containing any released CPS were saved, and the membrane pellets were suspended in water at a volume equal to the supernatant. As a negative control, an equal volume of water was added in place of the PLD.

### Alkaline treatment

Alkaline treatment was used for deacylation and was combined where indicated with NaBH_4_ to prevent sugar degradation by the peeling reaction (46–48). Mild alkaline treatment (aqueous base) was performed in 500-μl reactions containing 250-μl membranes isolated as above and NaOH at a final concentration of 0.1 N. Reactions were incubated for 30 min at room temperature, then neutralized with HCl and centrifuged at 135,000 × *g* for 30 min at 4°C to pellet the membranes, which were then suspended in water at a volume equal to the supernatant. Stronger treatment was performed with modification of the reaction mixture to contain 60% methanol v/v and is referred to as the methanolic base treatment. Supernatants to be used for GC-MS analysis were desalted using a Biogel P-2 column. Alkaline treatment with NaBH_4_ was performed under the aqueous conditions above with the addition of 10 mg/ml NaBH_4_. The reactions were then centrifuged at 135,000 × *g* for 30 min at 4°C. to pellet the membranes. The supernatant was neutralized with acetic acid in methanol (1:9, v/v) added dropwise until bubbling ceased. The membranes were suspended in water at a volume equal to the supernatant.

### Isolation of lipid-linked CPS

The D39 isogenic derivative Δ*cpsABCD* strain was used for isolating lipid-linked CPS, as this strain produces only short-chain CPS. As in the parent strain, the CPS is linked to both the peptidoglycan and membrane (16) (Gupta, Ambrose, Larson, and Yother, manuscript in preparation). Use of shorter chains improves the ability to detect non-CPS components during composition analysis. The Bligh-Dyer method (50) was used to extract CPS-lipid from one-ml of membranes isolated as above. The CPS-lipid was contained in the resultant aqueous-methanol fraction due to the high proportion of CPS relative to lipid. This fraction was applied to a 0.6 × 2.5-cm DEAE-Sephadex column equilibrated with 4 ml 100 mM ammonium acetate (pH 7.4) in 1:1 methanol:water and washed with 20 ml 1:1 methanol:water prior to sample application. The sample was applied in 0.5 ml 1:1 methanol water. The column was washed with 10 volumes of 1:1 methanol:water solution and CPS was eluted with a step gradient of 50, 100, 250, 500, and 1000 mM ammonium acetate (pH 7.4) in 1:1 methanol:water. Fractions were dried *in vacuo* and washed multiple times in methanol with subsequent drying *in vacuo* to remove the ammonium acetate. Each fraction was analyzed by GC-MS to identify the CPS-containing fraction, as described under Mass Spectrometry.

### Mass Spectrometry

Composition analyses by GC-MS were performed essentially as described previously (20). Samples for analysis were dried using vacuum centrifugation and subjected to 3 N methanolic HCl at 80°C for 16 hours in sealed glass ampoules. These samples were then dried *in vacuo*, and washed several times with methanol with drying *in vacuo*. The samples were transferred to a conical insert vial and trimethylsilated using 100 μl Tri-Sil HTP (Pierce) under argon for 1 hour at 80⁰C. Samples were analyzed on a GC–MS (Varian 4000, Agilent Technologies) fitted with a 30 m (0.25 mm internal diameter) VF-5ms capillary column. Column temperature was maintained at 160°C for 3 min and then increased to 260°C at 3°C/min and finally held at 260°C for 2 min, with a constant flow of 1.2 ml/min carrier gas. The effluent was analyzed by mass spectrometry (MS) using the electron impact ionization mode. Sugar identities and ratios were determined using purified sugar standards.

### Capsule immunoblotting

Fractionation of *S. pneumoniae* cell wall and membrane-containing protoplast fractions and analysis of CPS and LTA by immunoblotting were performed as previously described (16, 17).

## ACKNOWLEDGMENTS

This work was supported by NIH Awards R01 AI28457 (J.Y.) and T32 AI07041 (T.R.L.), and the Mizutani Foundation for Glycoscience (J.Y.). We thank Jan Novak for helpful suggestions regarding the work and manuscript.

